# Irregularity of instantaneous gamma frequency in the motor control network characterize visuomotor and proprioceptive information processing

**DOI:** 10.1101/2023.07.28.551050

**Authors:** Jihye Ryu, Jeong-woo Choi, Soroush Niketeghad, Elizabeth B. Torres, Nader Pouratian

**Affiliations:** Department of Neurosurgery, David Geffen School of Medicine at UCLA, Los Angeles, CA 90095, USA; Department of Neurological Surgery, UT Southwestern Medical Center, Dallas, TX 75390, USA; Psychology Department, Rutgers University Center for Cognitive Science, Computational Biomedicine Imaging and Modeling Center at Computer Science Department, Rutgers University, Piscataway, NJ 08854

**Keywords:** Goal-directed movement, reaching, instantaneous gamma frequency, entropy, motor network, electrocorticography (ECoG)

## Abstract

**Background:** Goal-directed movements involve integrating proprioceptive and visuo-motor information. Although the neural correlates of such information processing are known, the details of how sensory-motor integration occurs are still largely unknown.

**Objective:** The study aims to characterize movements with different sensory goals, by contrasting the neural activity involved in processing proprioceptive and visuo-motor information. To accomplish this, we have developed a new methodology that utilizes the irregularity of the instantaneous gamma frequency parameter for characterization.

**Approach:** In this study, 8 essential tremor patients undergoing an awake deep brain stimulation (DBS) implantation surgery repetitively touched the clinician’s finger (forward visually-guided/FV movement) and then one’s own chin (backward proprioceptively-guided/BP movement). Neural electrocorticographic (ECoG) recordings from the motor (M1), somatosensory (S1), and posterior parietal cortex (PPC) were obtained and band-pass filtered in the gamma range (30-80Hz). The irregularity of the inter-event intervals (IEI; inverse of instantaneous gamma frequency) were examined as: 1) correlation between the amplitude and its proceeding IEI, and 2) auto-information of the IEI time series. We further explored the network connectivity after segmenting the FV and BP movements by periods of accelerating and decelerating forces, and applying the IEI parameter to transfer entropy methods.

**Results:** Conceptualizing that the irregularity in IEI reflects active new information processing, we found the highest irregularity in M1 during BP movement, highest in PPC during FV movement, and the lowest during rest at all sites. Also, connectivity was the strongest from S1 to M1 and from S1 to PPC during FV movement with accelerating force and weakest during rest.

**Significance:** We introduce a novel methodology that utilize the instantaneous gamma frequency (i.e., IEI) parameter in characterizing goal-oriented movements with different sensory goals, and demonstrate its use to inform the directional connectivity within the motor cortical network. This method successfully characterizes different movement types, while providing interpretations to the sensory-motor integration processes.

## Introduction

Reaching to press an elevator button (visual goal) and reaching to scratch one’s face (proprioceptive goal) are movements that involve different sensory-motor processes. Although both biomechanical movements engage the arm’s joints and end effector (the hand) to accomplish the end goal, the brain must process these movements differently, because each requires different sensory processes (1–3) and force dynamics (i.e., when to flex and extend the joint muscles) (4–7). For that reason, it is likely that these distinct movements would be differentiated physiologically at the cortical level. However, there is a lack of methodology of using the cortical electrophysiological signals to characterize and differentiate these movements that are guided by different sensory goals.

A framework that explores the dynamics of goal-directed movements within the context of efferent and afferent streams of information flow is the internal forward models of action (e.g., principle of reafference (8), internal forward model (9,10)). In its current version, the model posits that when planning a movement, a motor command is sent down the spinal cord, and a duplicate motor command (termed efference copy) is sent to the posterior parietal cortex (PPC) to predict the afferent consequence of one’s self-generated movement, thus allowing for a faster and precise control. Nevertheless, the details of how sensory motor integration is made to execute the self-generated movement in the context of these models are still largely unknown. This is mainly because characterization of neural activities associated with various goal-directed movement have not been made dynamically, but instead commonly resorted to examining the grand-averaged epochs and applied assumptions of stationarity and linearity. In fact, conventional methods like the power spectrum fails to distinguish movements with different sensory goals (as shown in S1 Figure). Although such methods with simplifying assumptions help to reveal certain aspects of the motor network (e.g., decreased beta and increased gamma power during movement in (11–13)), we argue that utilizing a set of dynamical, nonstationary, and nonlinear analytical methods can capture the finer details that are hidden in the moment-to-moment variability (14), and therefore help to understand the mechanism behind sensory integration and movement planning.

In this study, we aimed to characterize the neural activities associated with goal-directed movements that involve different sensory goals. Here, we had patients with essential tremor undergoing an awake deep brain stimulation (DBS) implantation surgery to perform a task of touching the clinician’s finger, and then one’s own chin repeatedly (Figure 1A). We conceptualize that the forward reaching movement of touching the clinician’s finger (visual goal) would require more visuo-motor (VM) information processing, and backward movements of touching one’s chin (proprioceptive goal) would require more proprioceptive information processing. During this, electrophysiological signals were obtained using electrocorticography (ECoG) at the motor (M1), somatosensory (S1), and PPC (Figure 1B).

**Figure 1.**
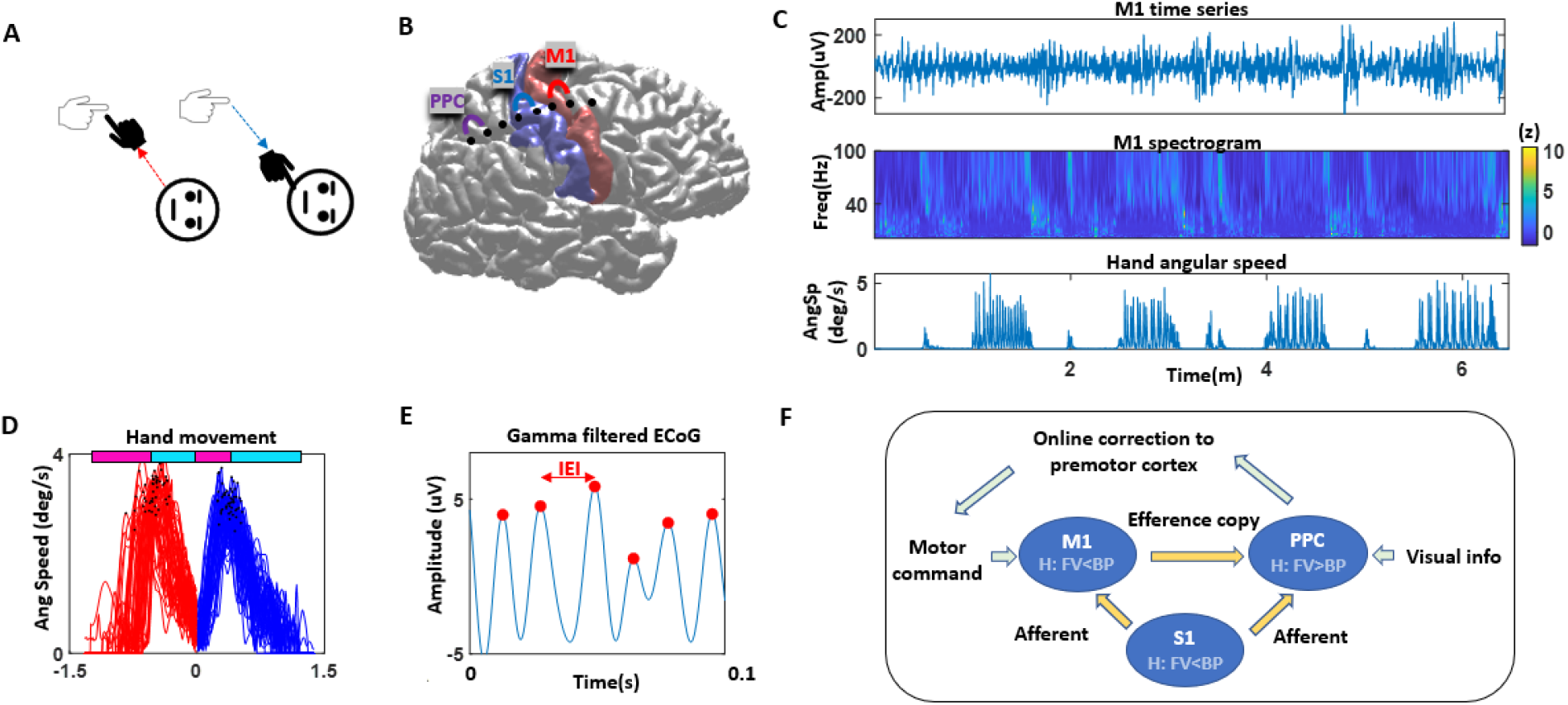
Experiment setup and analytics overview. **A.** Patient participant repeatedly performed a forward movement of touching the clinician’s finger, and then a backward movement of touching one’s own chin. **B.** ECoG strip was temporarily placed to cover the motor cortex (M1), somatosensory cortex (S1) and posterior parietal cortex (PPC). Channels were bipolar referenced to capture the local activity of the corresponding cortex. **C.** Representative participant’s electrophysiological signal at M1 (top) and its spectrogram (middle) and synchronized hand movement speed. **D.** Angular speed of the hand during forward visually-guided (FV; red) and backward proprioceptively-guided (BP; blue) movements, where time 0 is when the participant’s hand reaches the clinician’s finger. The pink horizontal line represents the time when the hand is applying accelerating force, and cyan line represents time when the hand is applying decelerating force. **E.** Electrophysiological time series data are gamma band filtered (30-80Hz), and the time between the maxima (denoted in red) is a parameter of the instantaneous gamma frequency. We refer to this as the inter-event-interval (IEI). **F.** Plausible model of the motor network in the context of the forward model of motor control. In this study, we characterized the neural activities associated with different movement types using the irregularity of the dynamical IEI at M1, S1, PPC. We also demonstrated how the transfer entropy of dynamical IEI’s can be applied to inform the directional flow of information (denoted in orange arrows) during different movement types.

To characterize the neural activities during movements with different sensory goals, we extracted the electrophysiological time series of when the participant performed a series of reaching tasks (Figure 1C), and compiled approximately 40 instances per participant (M = 41.1; SD = 11.9) of making forward visually-guided (FV) movements (i.e., touching the clinician’s finger) and backward proprioceptively-guided (BP) movements (i.e., touching one’s own chin) (Figure 1D). To further explore movements with different force dynamics, we also segmented these into times when the patient was accelerating force (i.e., when speed is increasing to its maximum) and when the patient was decelerating force (when speed is decreasing to zero). From the segmented dataset, we examined the instantaneous gamma frequency, which is quantified by taking the time between peaks within a gamma band filtered signal (termed inter-event-interval; IEI), as it is an inverse of the corresponding gamma frequency cycle (Figure 1E). The use of this parameter was first introduced in rodent hippocampus signals (15), demonstrating that the instantaneous gamma frequency reflects a nonlinear interplay between neural excitation and inhibition of interneurons.

Here, we compared the irregularity (i.e., less predictive) of the IEI during goal-oriented movements with a visual goal and a proprioceptive goal, and during a resting state, at the following cortical areas -M1, S1, PPC. Given that the gamma band power increases during movement (e.g., (11,12)), we hypothesize that active movement-related information processing would be characterized within the gamma frequency band, and that the magnitude of IEI irregularity would characterize the extent of such information processing. Specifically, we hypothesize the following (Figure 1F) - 1) M1 serves a critical role in executing movements (16), but also in integrating proprioceptive information (17). Given the dual role of movement execution and proprioceptive information processing, we hypothesize that M1 would show more irregularity during proprioceptive goal-oriented movement (i.e., BP movement) than a visual one (i.e., FV movement). 2) Since the S1 serves an important role in processing cutaneous and proprioceptive information processing (18), and given its strong connection to M1 (19), we hypothesize that it would exhibit a similar pattern as M1, but with a larger difference in irregularity between proprioceptive (BP) and visuo-motor information (FV) processes. 3) PPC has been well known to integrate visuo-motor (VM) information (20–22) at the intersection of motor and visual cortices. For that reason, we hypothesize that higher irregularity would be found during a visual goal-oriented movement (FV) than during a proprioceptive-goal oriented movement (BP). In addition, for exploratory purpose, we demonstrate the use of applying the IEI’s to transfer entropy (TE) metrics to understand the directional flow of information during goal-oriented movements (Figure 1F). Specifically, we show how the TE values vary across movements involving different sensory goals and force dynamics.

We highlight that this is a novel computational method of using the instantaneous gamma frequency (i.e., dynamical IEI) parameter in human ECoG signals to characterize neural activities associated with movement types with different sensory goals and force dynamics. This is possible because we relax the conventional assumptions of stationarity and linearity, and instead harness the moment-to-moment fluctuations of electrophysiological signals. Note, compared to conventional machine-learning methods that characterize movements (e.g., (23–25)), the strength of this novel approach is that it provides interpretations of neural activity, which we demonstrate with transfer entropy applications in this study.

## Materials and Methods

### Participants

Eight patients with essential tremor (demographics in Table S1), undergoing a bilateral or unilateral implantation of deep brain stimulation (DBS) leads targeting the ventral-intermediate nucleus (ViM) of the thalamus, were included in this study. All participants provided written consent approved by the institutional review board at the University of California, Los Angeles.

### Behavioral Task

In a single block, each participant was lying on the surgical bed, and was asked to raise the hand and posture for 30 seconds. Then, the participant was asked to touch the clinicians’ finger located within an arm-length, and then to touch one’s own chin repeatedly 10 times in a self-paced manner (FV angular speed Mdn = 1.6 deg/s; BP angular speed Mdn = 1.8 deg/s; details in Figure 3A) (Figure 1A). Subsequently the participant rested for 30 seconds, then started another block. 6 participants performed 4 blocks, and 2 participants performed 5 blocks in total. The entire study took approximately 7 minutes (Figure 1C). From here on, we refer to the movement of touching the clinician’s finger as a forward visually-guided (FV) movement, as it requires a visual goal, and the movement of touching one’s own chin as a backward proprioceptively-guided (BP) movement, as it requires a proprioceptive goal.

Here, we assume that the FV movement prioritizes visuo-motor information processing over proprioception, as they require attention to the visual goal, and the BP movement prioritizes proprioceptive information, as they require attention to the proprioceptive goal. Hence, when we mention “more information is involved”, we are referring to the relative priority/attention to the corresponding sensory information.

### Surgery and data acquisition

The recordings were made intraoperatively in an awake DBS surgery, during which an ECoG strip (8 channels with 1cm spacing; AdTech Medical, USA) was temporarily inserted subdurally via the burr hole made for DBS implantation for the purpose of research (26–28). For patients targeting the ventral intermediate (ViM) of the thalamus bilaterally, the ECoG strip was implanted through the right frontal burr hole; and for those targeting unilaterally, the ECoG strip was placed through the ipsilateral side of the burr hole. The burr hole was located at or approximately 1 cm anterior to the coronal suture (3 to 5 cm anterior to the central sulcus), and the ECoG strip was inserted posteriorly to cover the central sulcus. After all DBS leads were implanted, a lateral/sagittal fluoroscopy image was acquired, which showed the location of the ECoG strip along with the DBS leads.

For all participants, local field potentials (LFP) at the ViM of the thalamus (DBS target) and ECoG signals at the M1, S1, and PPC were recorded using a Matlab/Simulink software connected to an amplifier (g.Tec, g.USBamp 2.0). The signals were sampled at 4800Hz, and applied with a built-in 0.1Hz high band-pass and 60Hz notch filter. The ground and reference signals were obtained with a scalp needle inserted near the burr hole. For the purpose of this study, we analyzed the cortical signals obtained from the ECoG.

The participants’ kinematic signals were acquired with 2 opal inertial measurement unit (IMU) movement sensors (APDM, USA), then registered and sampled at 128Hz with the Motion Studio software (APDM, USA). The sensors were strapped on the moving hand and wrist. Here, we examined the wrist sensor’s angular velocity (deg/s) to extract the timing of differing movements (S2 Figure). Note, for the purpose of extracting the timing, we chose the wrist sensor (as opposed to the hand), as it is less susceptible to tremor. To temporally co-register the electrophysiological and kinematic signals, we used an external synchronization equipment (APDM, USA) that sent a digital output trigger to the electrophysiological signal amplifier to indicate the timing of the start and end of recording.

### Analysis

#### Preprocessing

With the aim of analyzing the motor (M1), somatosensory (S1), and posterior parietal cortex (PPC), we needed to anatomically localize the ECoG strip that was temporarily inserted during the surgery. To do this, we combined the pre-and post-operative CT scans and co-registered to the preoperative structural MRI, along with the lateral fluoroscopy image that showed the ECoG strip and implanted DBS leads. This method is adopted from (29), and details are further elaborated in our prior publications (30,31). Based on the visualized localization (Figure 1B), we identified the 2 closest channels that overlay the three cortical areas - M1, S1, and PPC - and bipolar-referenced those signals to capture the local activity in the three regions.

We visually inspected for electrical artefacts that showed clear evidence of artifact, based on an acute change in amplitude lasting more than 2 seconds, but did not find such segments. We also examined the power spectrum using the BOSC algorithm (32) at individual frequencies from 1 to 100Hz with 1Hz step width, and 6^th^ order wavelets. The power time series were normalized by z-scoring each frequency over the entire recording (approximately 7 minutes) per cortical site (Figure 1C; S1 Figure). Here, we confirmed an increase in gamma power (>30Hz) during movement for all participants at the motor cortex, consistent with prior findings (e.g., (11,28)).

In order to segment data for the times when the participant is performing a forward visually-guided (FV) movement, backward proprioceptively-guided (BP) movement, and at rest, we extracted the angular velocity of the wrist sensor, and computed the Euclidean norm to obtain a scalar angular speed. The angular speed profile informed us of when the participant reached the clinician’s finger, or one’s own chin, because the speed was near 0 (deg/s) at those times. These would define the timing of the start and end of either the FV or BP movement. The angular velocity along the z-axis informed us of the target that the participant has reached when the angular speed was near 0 (Figure 1D; S2 Figure).

Specifically, the direction of the angular velocity along the z-axis at zero-crossing points would inform us of the target. If the angular velocity changes from negative to positive, it has reached the chin, and when it changes from positive to negative it has reached the clinician’s finger. For some patients, their hand hovered around the clinician’s finger during the FV movements to precisely reach the finger for about 1 second or less. In these cases, we excluded those short moments of hovering, as this may be due to tremor and/or the clinician’s inadvertent moving (S2 Figure). To obtain the data during the resting period, we extracted the resting datasets (where angular speed is continuously near 0), and truncated the first 5 and last 5 seconds of the resting period within each block, to avoid any effect from movement preparation and/or change.

We separated the M1, S1, and PPC time series by movement types - FV movement, BP movement, resting - and then gamma band-pass filtered the signal (30-80Hz, 200^th^ order zero-phase, transition width 0.2, FIR). From the band-passed signal, maxima were identified (Figure 2A), and the time between two sequential maxima were defined as the inter-event intervals (IEI). In the following sections of analytics, we introduce three novel analytics that utilize this IEI parameter to characterize the different types of goal-oriented movements.

**Figure 2.**
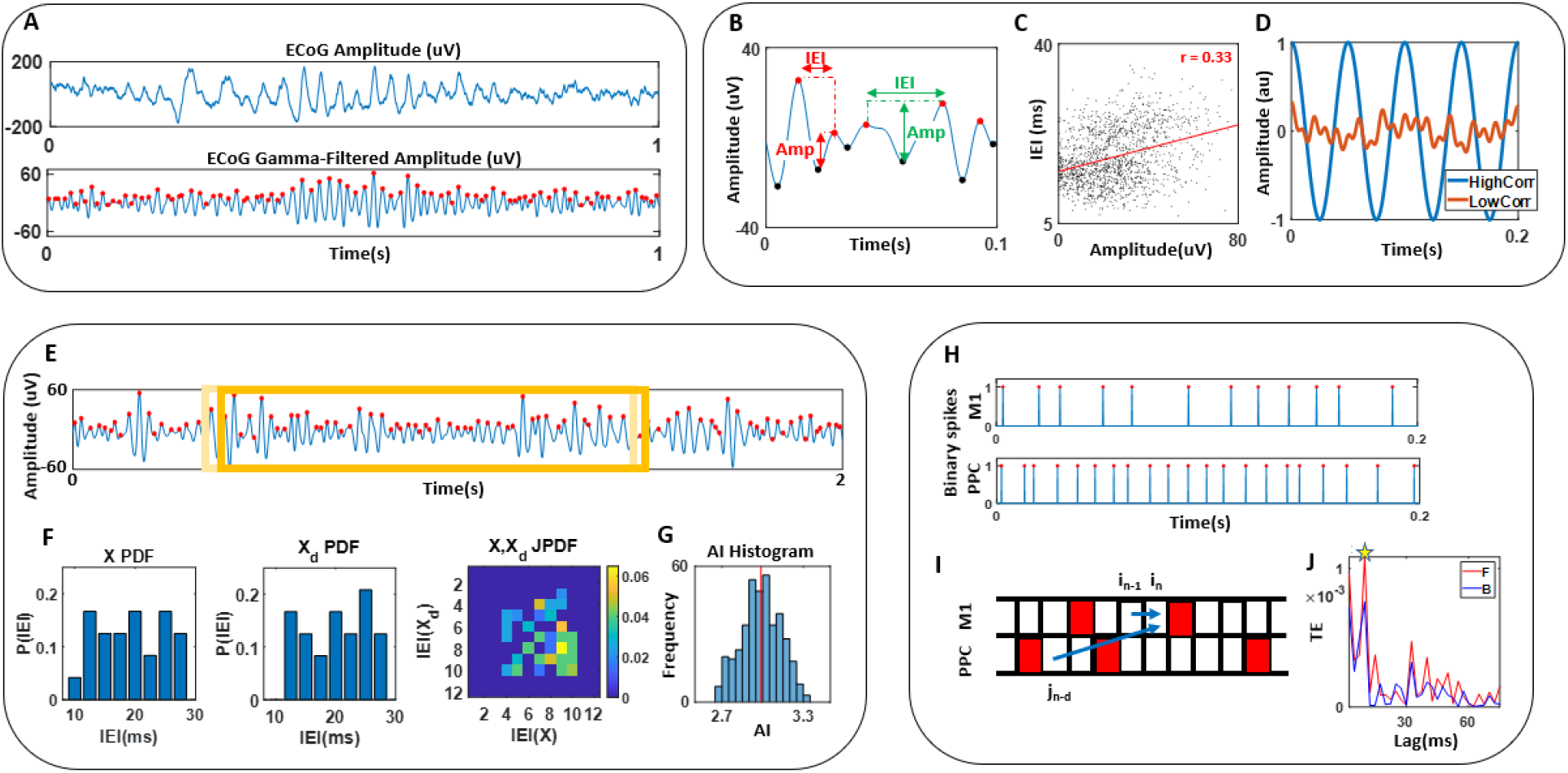
Analytics pipeline. **A.** ECoG signals were gamma band-pass filtered (30-80Hz) and the maxima were identified to compute the inter-event-interval (IEI). **B.** As a first set of analytics, IEI and its corresponding amplitudes were extracted. **C.** Correlation was computed for the paired dataset of IEI and amplitudes, for all movement types, per participant. **D.** We conceptualized that high correlation from **C.** exhibits a largely synchronized signal where the IEI and amplitude values are predictive. Conversely, low correlation would exhibit a more random irregular signal. **E.** As a second set of analytics, within 1-second time window, the IEIs were extracted and compiled (X), and this window was shifted by 5ms and again, IEIs were extracted and compiled (Xd). **F.** The PDF of X and Xd obtained from **E.** along with its JPDF were plotted, to compute the mutual information to characterize irregularity in the signal. We term this auto-information (AI). **G.** AI values were compiled and the means were extracted per movement types and per participant. **H.** As a last set of analytics, where we aim to see the directionality between M1, S1, and PPC, we created a binary spike train, where the maxima from **A**. are assigned 1, and the rest assigned as 0. **I.** Transfer entropy (TE) was computed pair-wise to assess the directional connectivity between two cortical areas, with lags varying from 2.5ms to 75ms. **J.** Maximal TE value was extracted within the varying lag values, per movement type (by sensory goals and force dynamics), and per participant.

#### Correlation between amplitude and IEI

Within a gamma band-pass filtered time series waveform, we identified a series of gamma cycles (i.e., valley with two maxima and one minimum in between) to extract the amplitude and IEI, where the amplitude is defined as the difference between the minimum and the subsequent maximum (Figure 2B). We compiled a set of paired data, comprised of amplitudes and the corresponding IEIs, and computed the correlation between the two (Figure 2C). For all participants, each movement types yielded approximately 2500 pairs of data to compute the correlation. Prior literature has shown positive correlation between the amplitude and IEIs within gamma cycles (15,33,34), and a recent study demonstrated how such positive correlation is expected in waveforms that contain a 1/f structure (34). Indeed, for all participants, statistically significant (p<0.01) positive correlation was found in all movement types within the range of 0.1 to 0.45 in all cortical areas.

We aimed to compare the magnitude of correlation across different movement types, using a paired Wilcoxon signed-rank test across all 8 participants. Here, we assume that largely synchronized signals are highly correlated, and asynchronous signals are less correlated, and that low correlation characterizes new information actively being processed (Figure 2D). Note, the gamma band power is not sufficient to differentiate and characterize the FV and BP movements, as shown in S1 Figure.

#### Auto-information across IEIs

Another way to characterize movements with a visual goal and a proprioceptive goal, is to compute the mutual information (based on information theory; (35)) between two sequential time windows from a single time series. We refer to this as auto-information (AI) (conceptually similar to auto-correlation). Specifically, we down-sampled the data from 4800Hz to 400Hz, and concatenated all trials within the same movement type into a single time series. These were then gamma band-pass filtered, yielding a series of IEI’s (Figure 2E). Per 1 second time window, these IEIs were plotted on a discrete probability distribution function (PDF) with a range spanning from 2 frames (5ms = 2 / 400hz) to 14 frames (35ms = 14/400hz). This 1 second time window was then shifted forward by 2 frames (i.e., 5ms), and IEI’s from this window were plotted again on a PDF. Then, with the same bin size of 1 frame (2.5ms), a joint PDF between these two time windows were plotted (Figure 2F). We chose to down-sample the data to 400hz, because the minimum IEI was found to be 2.5ms based on the 4800hz gamma filtered data, and the duration of a single frame within a 400hz data equals 2.5ms. Also, we chose the time window length as 1 second, because this was the shortest time length to gather sufficient IEI datapoints (approximately 100) to produce a meaningful frequency distribution. We chose the shift size of 2 frames (5ms) since this would be a short enough time shift to capture the changing IEIs (as the shortest IEI is 2.5ms). Given the obtained two single probability distributions, and the corresponding joint probability distribution, we computed the auto-information (i.e., mutual information) with the following formula (35):

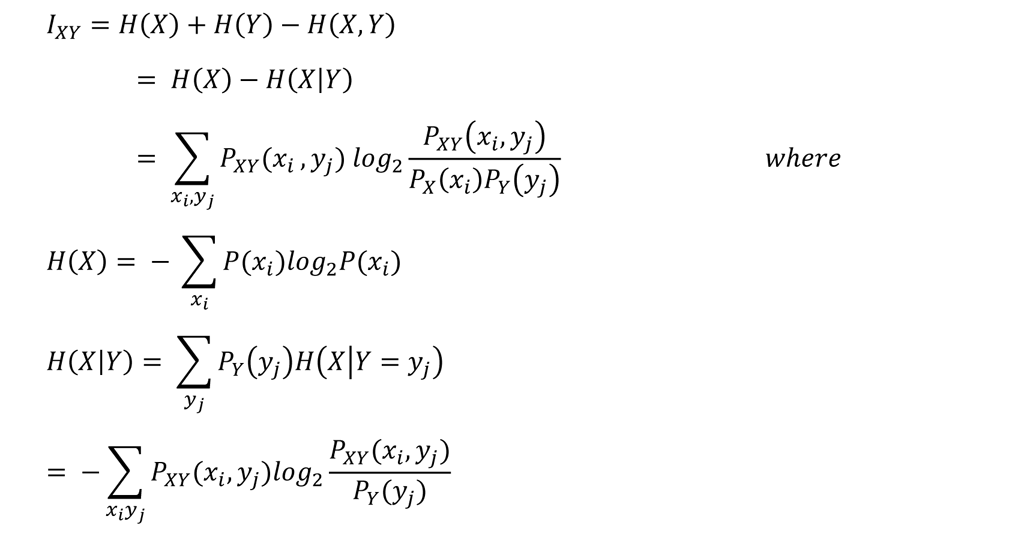

Function *H* is an entropy function, *X* is the IEI variables from the first time window, *Y* is the IEI variables from the subsequent (shifted) time window, *i* and *j* are IEI bins ranging from 2 (frames) to 14 (frames). The details of this mathematical derivation in the context of neuroscience is well explained in (36). Overall, this AI metric represents a stochastic dependency between the two sets of variables *X* and *Y*. If the stochasticity of the two sequential time windows were highly dependent on each other, they would yield a high AI value; and if the two were entirely independent, the AI value would equal 0.

We computed the AI (i.e., mutual information between the two sequential time windows shifted by 5ms), then shifted these two sets of windows by 10% (100ms) and computed the AI values again. This resulted with approximately 400 AI values per movement type for each participant. For comparison, we took the mean of these approximately 400 AI values (Figure 2G), then compared the means between the different movement types using a paired Wilcoxon signed-rank test across all 8 participants.

#### Transfer entropy

Lastly, we introduce how the instantaneous gamma frequency (i.e., dynamical IEI) can be applied to understand the directionality of pairwise informational flow within the three cortical areas (M1, S1, PPC) during different movement types. Note, for the purpose of exploration, we separated the datasets by sensory goals (i.e., FV, BP, rest), and further segmented them by force dynamics. That is, for each FV and BP movement, these were segmented by times when force was accelerating (time when angular speed changes from zero to its maximum) and when the force was decelerating (time when angular speed changes from its maximum to zero) (Figure 1D).

With these segmented datasets, we created a binary spike train for each movement types, and computed the transfer entropy (TE) between each pair of cortical areas. Specifically, we down-sampled the data from 4800Hz to 400Hz, concatenated all trials within the same movement type into a single time series (amounting to approximately 45 seconds per movement type), then gamma band-pass filtered the data. Then we identified the indices when the peaks occurred, and created a binary spike train where the maxima indices were assigned the value 1, and the rest were set to 0 (Figure 2H). Per a single movement type time series data (with an approximate total length of 45 seconds), we found approximately 2500 spikes. These were then used to compute a set of delayed TE using the toolbox developed by (37), where the following formula was used to compute the transfer entropy of *J* preceding *I* with *d* delay :

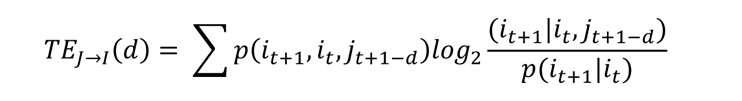

*J*and *I* corresponds to 2 cortical areas, *i*_*t*_ is the binary value at time *t* (i.e., frame *t*), and *d* is the delay period. This metric essentially measures how much prediction of *I* is improved, when we know the past values of *I* from 1 frame (2.5ms) ago and *J* from *d* frames ago, as opposed to knowing just *I* from 1 frame (2.5ms) ago (Figure 2I). Here, we examined the TE values at delay periods 1 to 30 frames (i.e., 2.5ms to 75ms) in 1 frame (2.5ms) increment, and extracted the maximal TE value within such range of delay period (Figure 2J).

We compiled these maximal TE values per movement types, for each participant. We then compared across movement types using a paired Wilcoxon signed-rank test across all 8 participants for the following directions – M1 to PPC, S1 to PPC, and S1 to M1.

## Results

As a first step, we compared the mean angular speed of the hand during the forward visually-guided (FV) (M = 1.94, n=8) and backward proprioceptively-guided (BP) (M = 1.89, n=8) movements, and confirmed that they were not different (p=0.92) (Figure 3A). In addition, we examined the mean IEIs during the three movement types – FV, BP, Rest - and found the values to be highest during rest in all cortical areas (Figure 3B-D). Furthermore, IEIs were lowest during BP movement in M1, and lowest during FV movement in PPC. Although such findings may imply that the mean IEIs may be a sufficient metric to characterize the three movement types, we found the frequency histogram of the IEIs in M1 and PPC to roughly exhibit a bimodal distribution (Figure 3E-F). Note, the Hartigan’s dip significance test of the distributions’ unimodality had shown p-values ranging from 0.03 to 0.08. This bimodality is due to an artefact of applying the 60hz notch filter, which was an inevitable limitation to the study environment. For that reason, given the bimodality, we deem the mean IEI values to be a limiting metric for characterization, and thus rely on the dynamical changes in the IEIs to be more appropriate to characterize the different movement types.

**Figure 3.**
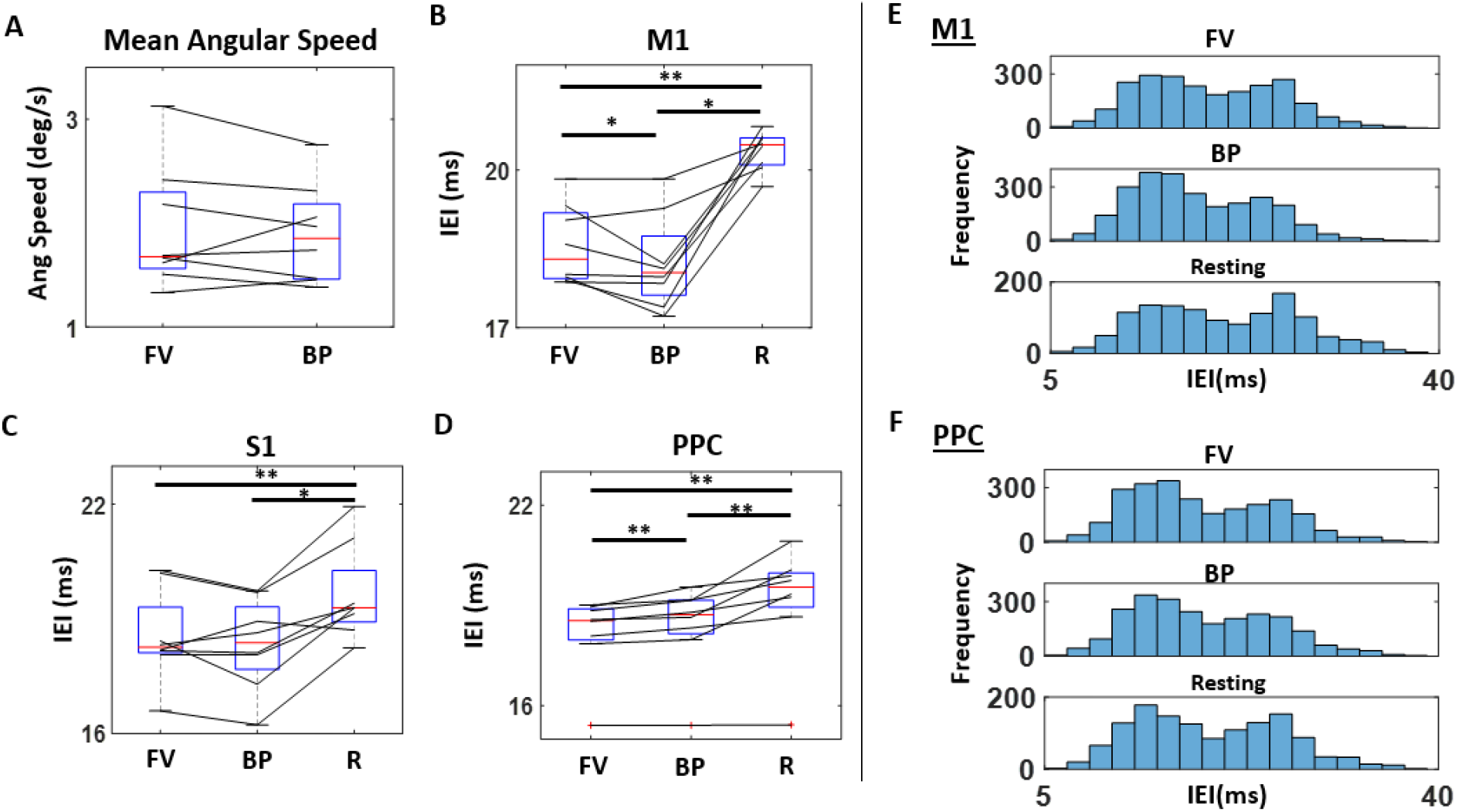
Mean angular speed and IEI. **A.** Mean angular speed during forward visually-guided (FV) and backward proprioceptively-guided (BP) movements for 8 participants. **B.** Mean IEI during FV and BP movement, and rest (R) for all 8 participants in M1, **C.** in S1, and **D.** in PPC. **E.** Frequency distribution of IEI’s from a representative participant during FV (top) and BP movement (middle) and rest (bottom) in M1, **F.** and in PPC. The distribution is slightly bimodal, indicating that the mean as a summary statistic is not optimal to characterize the differing movement types. *p<0.05, **p<0.01

### Low correlation of gamma cycle IEI and amplitudes characterize active new information processing

We correlated the amplitudes and corresponding IEIs per movement types (FV, BP, Rest) and per participant, and found a positive correlation in the range of 0.1 to 0.45 (all participant and movement type showed a significant correlation at p<0.01). We hypothesized that lower correlation (i.e., irregularity) in the amplitudes and IEIs would reflect active new information processing, such that M1 and S1 would show the lowest correlation during BP movement (i.e., proprioceptive information) and PPC during FV movement (i.e., visuo-motor information); and that highest correlation (i.e., most regularity) would be found during resting state in all cortical areas.

Assuming more proprioceptive information processing occurs in M1, as hypothesized, M1 showed a lower correlation during BP movement (M=0.31, n=8) than during FV movement (M=0.33, n=8) (p=0.04, paired Wilcoxon signed-rank test; Figure 4A). Also, assuming resting state to be associated with the least new information being processed, we found the resting state to exhibit the highest correlation (M=0.38, n=8). In S1, we did not find differences between FV (M=0.31, n=8) and BP (M=0.30, n=8, p=0.04) movements (p=0.3), but found the resting state (M=0.35, n=8, p=0.05) to exhibit higher correlation compared to FV (p=0.05) and BP (p=0.04) movements (Figure 4B). In the PPC, we did not find differences between movement types (FV vs. BP p=0.38; FV vs. Rest p=0.2; BP vs. Rest p=0.25). Overall, we find the resting state to have the highest correlation in all three cortical areas. This strengthens our hypothesis that irregularity in the gamma IEI is reflective of active new information processing. Note, we also visualized the gamma band filtered signals per movement type, but such patterns of correlations are not easily noticeable with the naked eye (Figure 4E).

**Figure 4.**
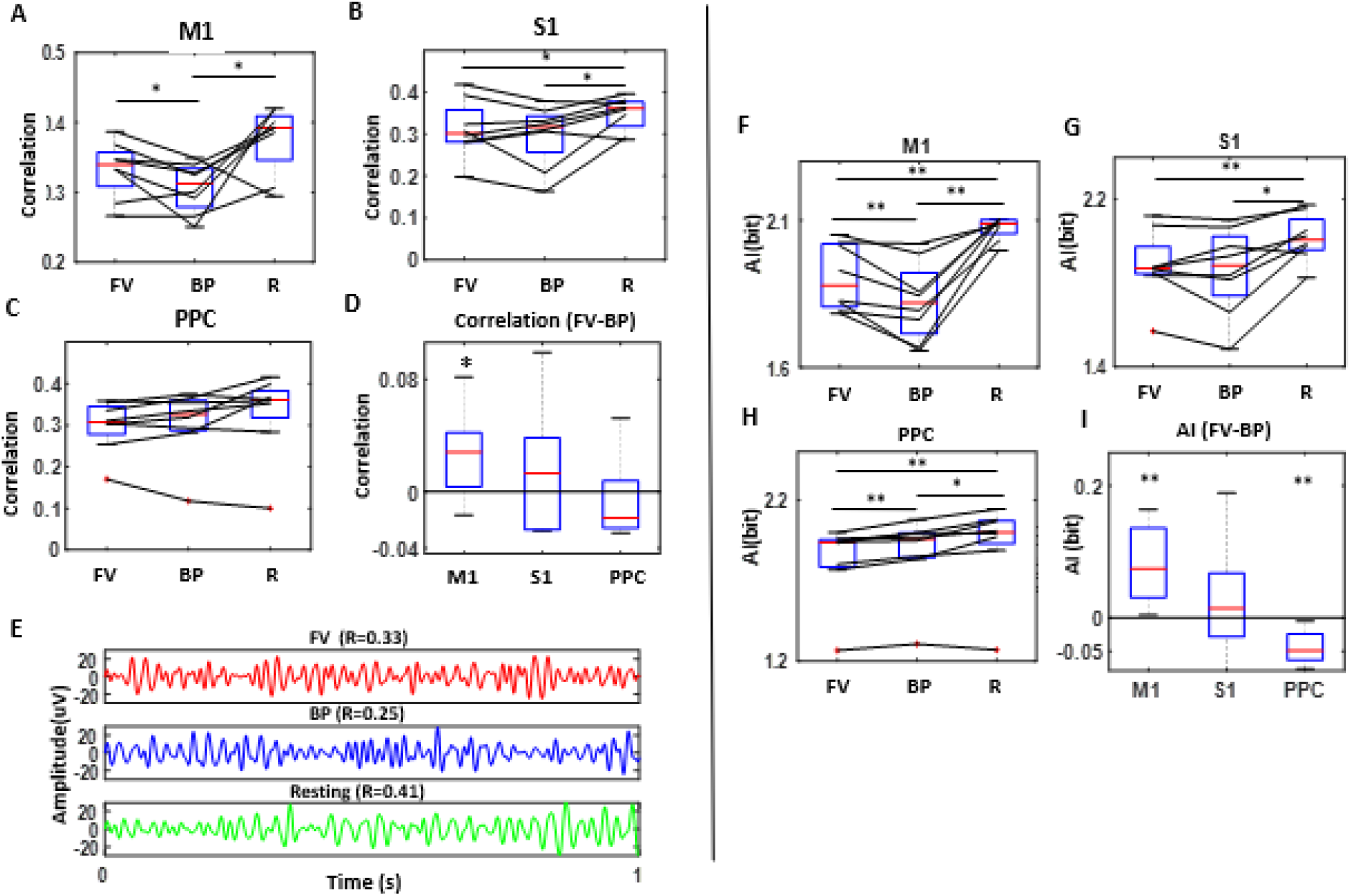
Irregularity of IEI represented by the correlation with its corresponding amplitude, and auto-information of IEI time series. **A.** Correlation between IEI and amplitudes are compared for all 8 participants between forward visually-guided (FV), and backward proprioceptively-guided (BP) movements and rest (R) in M1, **B.** S1, and **C.** in PPC. Generally, we find the resting state to have highest correlation. **D.** In distinguishing the FV and BP movements, M1 shows the largest difference. **E.** The variation in correlation across movement types are not easily visible by the naked eye. **F.** AI comparison for all 8 participants between forward visually-guided (FV), and backward proprioceptively-guided (BP) movements and rest (R) in M1, **G.** S1, and **H.** in PPC. **I.** FV and BP movements are most differentiable in M1 and PPC. *p<0.05, **p<0.01

### Low auto-information (AI) of gamma cycle IEIs characterize new information processing

As another way to characterize the irregularity in the signal to reflect new information processing, we computed a series of AI of the IEI stochasticity between two sequential 1s windows, that are shifted by 5ms. Low AI would indicate more irregularity in signals (i.e., more independence from the past) where new information is being processed, and high AI would imply a more regular signal (i.e., more dependence from the past).

In M1, assuming that BP movement would involve the most proprioceptive information to be processed, as hypothesized, BP movements showed the lowest AI (M=1.82, n=8), then the FV movement (M=1.90, n=8), and the highest AI value during resting state (M=2.07, n=8) (Figure 4F). On the other hand, in S1, the BP movement (M=1.86, n=8) did not show difference from FV movements (M=1.88, n=8), which is contrary to what we hypothesized (p=0.55). Still we found the highest AI during resting state (M=2.02, n=8) in S1, and resting state to be different from the two movement types (FV p<0.01; BP p=0.02) (Figure 4G). In PPC, assuming that FV movement would involve the most visuo-motor information to be processed, as hypothesized, FV movement showed the lowest AI (M=1.83, n=8), then the BP movement (M=1.87, n=8), and the highest AI during rest (M=1.93, n=8) (Figure 4H). We also computed the AI of two 1s windows that are shifted by 20ms (not 5ms), and also found a similar pattern with statistical significance as well (S3 Figure).

### Dynamical IEI inform the directionality connectivity between M1, S1, and PPC during movement

For exploratory purpose, we demonstrate how the dynamical IEI parameter applied to transfer entropy methods can inform the directional interactions between M1, S1, and PPC. Generally, we found the lowest connectivity between the three cortical areas during rest, and this confirmed that the metrics extracted from these areas indeed characterized movement-related activities (**Error! Reference source not found.**Figure 5A-C). Here, we also found a stronger directional flow during FV than BP movements from S1 to PPC (p=0.01), and from S1 to M1 (p=0.08) (Figure 5**Error! Reference source not found.**D). When we further segmented the movements by force dynamics, we observed the strongest connectivity when force was accelerated than when decelerated (Figure 5**Error! Reference source not found.**E-G), and the difference was most pronounced during FV movements.

**Figure 5.**
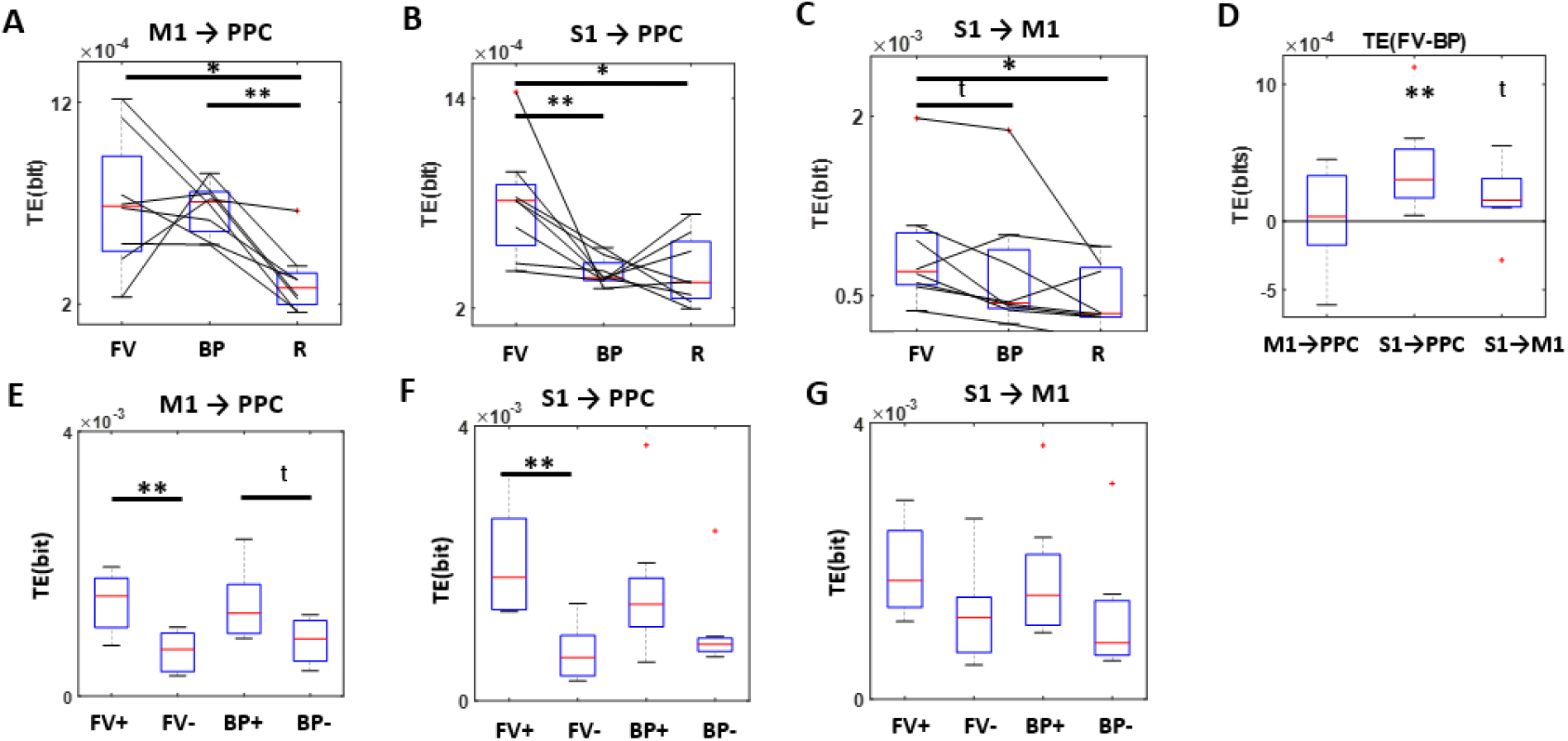
Directional connectivity of the dynamical IEI compared across movements with different sensory goals (top; A-D) and further segmented by force dynamics (bottom; E-G). **A.** Directional connectivity is assessed with transfer entropy from M1 to PPC, **B.** from S1 to PPC, and **C.** from S1 to M1. They all show lowest connectivity during rest, confirming that these represent movement-related interactions. **D.** The connectivity between S1 and PPC, and S1 and M1 is higher during FV than BP movements. **E.** Directional connectivity from M1 to PPC, F. from S1 to M1, and G. from S1 to M1, was examined by further segmenting movements by accelerating force (FV+, BP+) and decelerating force (FV-, BP-). Generally, connectivity is strongest when force is accelerated, and is most pronounced during FV movements. *p<0.05, **p<0.01, ^t^ p<0.1

Also, given the conceptualization of a directional flow from M1 to PPC during movement to represent an efference copy (based on the internal forward model of movement (8–10)), we do observe higher TE values during FV (M=7e^-4^, n=8) and BP movement (M=6.7e^-4^, n=8) than during rest (M=3e^-4^, n=8) (Figure 5**Error! Reference source not found.**A). Although there was no difference between FV and BP movements (p=0.7), we do find difference when force is accelerated than when decelerated during FV (p<0.01).

Lastly, we also examined other pairwise directional flow among the three cortical areas but did not find difference between the FV and BP movements (S4 Figure). However, when we further segmented the movements by force dynamics, we generally find a stronger connectivity during accelerating FV than decelerating FV movements – specifically from M1 to PPC (p<0.01) and PPC to M1 (p<0.01), from M1 to S1 (p=0.02), and from S1 to PPC (p<0.01) (S4 Figure). Such difference in force dynamics is muted during BP movements, and the only difference is found in the direction from M1 to S1 (p=0.02) (S4 Figure). We also examined the optimal lag values when TE was maximal for each participant, and found that the majority exhibited this lag to be at 2.5ms, and often at around 10ms (S5 Figure). Overall, we find the connectivity between M1, S1, and PPC to be higher during movement than rest, higher during FV than BP movements, and higher during accelerating than decelerating movements.

## Discussion

We demonstrate a novel methodology of characterizing goal-oriented movements with differing goal modality (i.e., visual versus proprioceptive goal), conceptualizing that the irregularity in IEI reflects active new information processing. We do this by harnessing the moment-to-moment variability in the gamma band-pass filtered ECoG signals, and thereby capturing the nonstationary and nonlinear features. We also show how the dynamical IEI changes can inform us of the directional connectivity by providing exploratory results. Specifically, we find the connectivity to be strongest during a visually guided goal-oriented movement (FV) with accelerating force (FV+), and weakest during rest. We also show preliminary empirical evidence of an efference copy using this parameter, as we find a strong connectivity from M1 to PPC during movement.

We emphasize that using a nonstationary parameter (i.e., gamma IEI) and its dynamical changes allow us to characterize the activity within a local cortical area, and can inform us of the interactions within the motor control network (i.e., M1, S1, PPC). Conventional ways of capturing information processing entail searching for an increased oscillatory power, but these do not reflect the dynamical and nonstationary features of the brain signals. Indeed, we show that a conventional power spectrum method fails to differentiate the two movement types (FV and BP) (shown in S1 Figure). With prior knowledge on the role of M1 and S1 in regards to proprioceptive information processing, and PPC on visuo-motor processing, we demonstrate that the irregularity in gamma IEI reflects such local activation of information processing, and captures the finer differences in sensory processing. This is possible because we are reflecting the dynamical and nonstationary features of cortical activity, by relaxing the stationarity and linearity assumptions while harnessing the moment-to-moment variability of the oscillatory ECoG signal. Such attempts are absent in the conventional epoch-based analyses, because these methods involve averaging out the moment-to-moment variability with the general assumption that the cortical electrophysiological signal follows a stationary Gaussian distribution. We also highlight that our novel method provides a way to overcome the inherent difficulty in assessing the gamma band signal which has a low signal-to-noise ratio. This is possible because we are extracting large amounts of data, and thus increasing the statistical power. For instance, the moment-to-moment variability in the IEIs yield in the magnitudes of 3000 data points per 1-minute recordings (3000 data points = 1000ms / Average IEI 20ms * 60 seconds). For such a short amount of time, our method provides a large dataset to analyze and thus compensate for the low signal-to-noise ratio. This is indeed a large merit compared to common characterization methods like machine-learning, which require a very long time of data collection and training. Moreover, the interpretability of our method (e.g., demonstrated by directional connectivity) provides an added benefit compared to the common machine-learning methods, as these do not provide much knowledge on the interactions that occur within the cortical network.

We interpret the irregularity of the gamma IEI to reflect the *relative attention* of new information processing. In fact, when we hypothesize that BP movement would involve more proprioceptive information processing in the M1, we assume that given a limited capacity of attention, proprioceptive information would be weighted (attended to) more than the VM information, and that the irregularity in M1 would reflect such difference (Figure 6A). Because FV and BP movements require a similar linear trajectory of hand movement (but (38)), we assume that the proprioceptive information itself would be similar between the two movements. For that reason, the irregularity in M1 signal would not reflect the amount of proprioceptive information processing per se, but rather how much attention is devoted towards that information processing.

**Figure 6.**
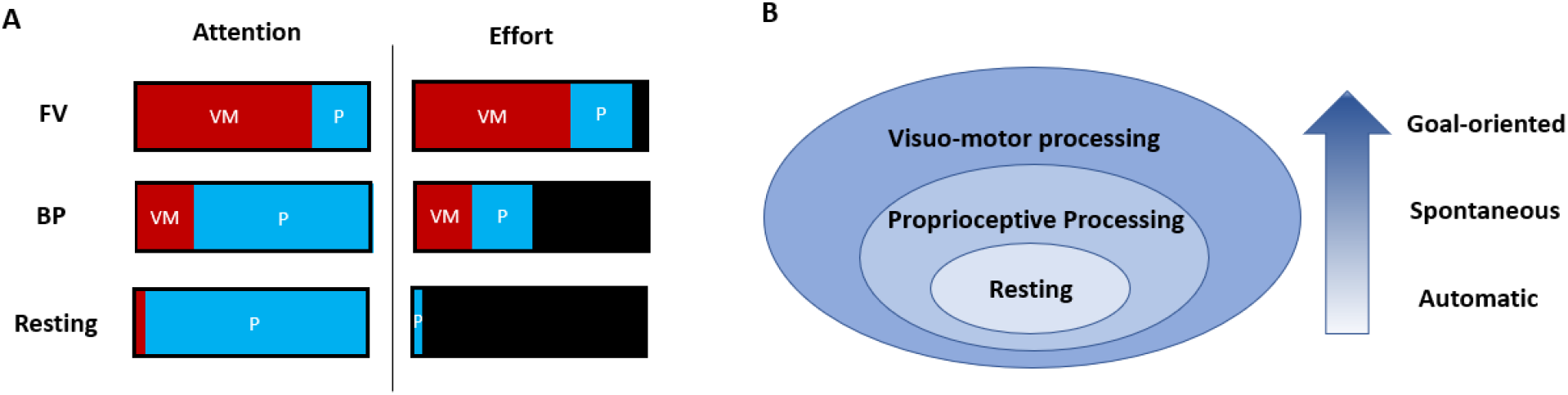
Schematic organization of the motor control network. **A.** Under the relative attention model (Attention), we conceptualize that the irregularity of M1 signal reflects the increased *attention to* proprioceptive information. We argue that higher irregularity in M1 during BP movement does not reflect a lower physical effort (Effort), because the irregularity is lowest during resting state and highest during BP movements, and physical effort is highest during FV movement. **B.** Stronger connectivity within the motor control network are found during FV movement (requiring more VM processing) than BP movement and the least connectivity during resting state. We also find stronger connectivity when force is accelerated than decelerated during both FV and BP movement. We speculate this level of connectivity to reflect an organization where the connections strengthen according to the developmental order of neuromotor control – where automatic functions develop first, and goal-oriented movements develop later, and movements exerting low-level force develop first, and those exerting more force to develop later.

Still, we provide alternative interpretations for the observed differences. One possibility is that the difference may reflect the level of physical effort to execute the two types of movement. This is because the FV movement required the participant to reach up against gravity, and BP movement to move towards gravity. This is a limitation of an intraoperative setting, where the awake patient had to lie supine, and was inevitably required to move up against gravity to reach a visual goal. If the difference in irregularity is due to such differing levels of physical effort, however, higher irregularity would mean involving less physical effort, as BP movements showed higher irregularity in M1 than FV movements. However, the irregularity was found to be the lowest during rest, implying that the level of irregularity is not likely to depend on the level of physical effort (Figure 6A).

Another possible reason for the difference in irregularity between FV and BP movements may simply be due to the different risks associated with the movement. That is, FV movement involves reaching for an external target and does not entail much risk to harm oneself, leading the movement to be more forceful. On the other hand, BP movement involves the risk of hitting oneself in the face, as this is moving against the natural momentum of gravity, and thus require more control and “braking” along the way. For that reason, the difference in irregularity may reflect the accelerating forces in movement, that is modulated by risks involved. However, when we compared the irregularity across movements involving different force dynamics (i.e., accelerated FV versus decelerated FV movement, accelerated BP versus decelerated BP movement, shown in S6 Figure), we did not see much difference. For that reason, we argue that the irregularity in gamma IEI would most likely reflect the general level of attention, rather than the force generated by the hand.

In order to demonstrate the use of the dynamical Gamma IEI parameter, we provided exploratory results of the motor network’s cortical connectivity using transfer entropy methods. Here we found the strongest connectivity during accelerated FV movement and the least during the resting state. This implies that the motor network strengthens its connection when it exerts a higher level of neuromotor control. Here we assume that the FV movement involves the highest level of neuromotor control (14,39), as it integrates information from the external world along with its internal body (i.e., both visuo-motor and proprioceptive information processing), whereas the BP movement mainly processes information from within the internal body. We also assume that accelerated movement involves more control, as a higher level of force is exerted than when it is decelerating. It is possible that such varying connectivity strength reflects the order of neuromotor development, because we speculate that the motor network is weakly connected at birth, when simple autonomic activities are mainly executed. As one matures and exerts more goal-directed movements that require a higher level of neuromotor control, the motor network would strengthen its connectivity (Figure 6B). Nevertheless, we caution that this varying level of connectivity strength may be due to the levels of physical effort. In fact, the stronger connectivity found during accelerated movements compared to decelerated movements support such reasoning. For that reason, in a follow up study, it will be helpful to have a control condition where a FV movement would not be physically effortful. This would be possible to do within a sitting environment, using an EEG or MEG.

We also found stronger connectivity from M1 to PPC during movement compared to rest, which is a potential empirical evidence of an efference copy from the forward models of motor control (e.g., reafference-cancelling model (8), internal forward model (9)). These models postulate that predictive codes are sent from M1 to PPC to forecast the resulting sensations of self-generated movements, thereby provide a better control of one’s movement. In the past, a common empirical evidence of an efference copy has been sensory attenuation (40–42) during active self-generated movement compared to passive movement. Here, we provide a different angle of evidence, where we directly show a stronger informational flow from M1 to PPC during self-generated movements. Interestingly, such connectivity does not distinguish the two movement types – FV and BP - implying that the efference copy may be indifferent to sensory goal modality. On the other hand, we do find a stronger mutual connectivity between M1 and PPC during accelerated FV movement compared to decelerated FV movement. We speculate that the efference copy may not distinguish the finer differences in sensory processing, but may instead distinguish the force generated by the end-effector. This may be why we see the difference during movement versus non-movement.

However, we warrant several limitations to these claims. The location of the PPC channel across participants were not consistent. Within the broad region of PPC, some participant’s PPC contacts were more superior (vs. inferior) and some were more posterior (vs. anterior). Given that the PPC has differing functional zones (22,43), among which includes postural information processing (44,45), it is indeed important to record from the precise functional region for reaching and consistently across participants. However, due to the limited timeframe during an awake DBS surgery, achieving a precise and consistent placement of the ECoG strip across patients entails clinical risk. Another limitation is that passive movements were not examined as a control condition. For that reason, it is possible that information flow from M1 to PPC is merely reflecting proprioceptive information processing that occurs during any movement, regardless of whether it is active or passive. If that is the case, this connectivity finding may not be a relevant evidence for an efference copy, but rather a simple explanation of how the brain detects movement. In a follow up study, it will be helpful to verify this by having a passive movement condition.

As a last point of discussion, contrary to our hypothesis, we find M1 to be more active in proprioceptive information processing (i.e., show larger difference in irregularity between BP and FV movement) than the S1. We originally hypothesized that S1 would show a larger difference between the two movements, as we assumed S1 to reflect more active proprioceptive processing. Although we know that both M1 and S1 are involved in processing proprioceptive information, the results indicate that M1 may be more active in processing such information than S1. We conjecture that since there’s a stronger flow of information from S1 to M1 during movement, perhaps S1 is a general receiving site for continuous bodily information, whereas M1 is where select information pertaining to movements are processed. Due to its selectivity, perhaps this is why there is a larger differentiation between the two (FV and BP) movement types.

We acknowledge that the small sample size of 8 participants and the clinical diagnosis of the sample (essential tremor) limits the generalizability of this study. These are factors inevitable to invasive recordings on human participants, as the population size that would undergo such invasive recording is small to begin with. However, we point out that a single data point summarized for each participant is based on a very large dataset (appx 2500 datapoints) from that person, and is thus a summary statistic with high statistical power on its own. Still, the data was obtained from essential tremor patients, who have impairment in movement, and we do not know how much of these can be generalized to the neurotypical population. To that end, it would be helpful to verify this with the neurotypical population using high density EEG in a follow-up study.

In summary, we introduce a novel methodology that utilize the instantaneous gamma frequency (i.e., Gamma IEI) parameter in characterizing goal-oriented movements with different sensory modality, and demonstrate its application to reveal the directional connectivity within the motor cortical network. This was possible because we relaxed the stationarity and linearity assumption, and captured the dynamical changes by harnessing the moment-to-moment variability from the oscillatory cortical signals. Through this method, we demonstrate how the irregularity in the gamma IEI informs the state of active new information processing, and how applications to transfer entropy methods can inform the directional connectivity within the motor network.

## Supporting Information

**S1 Table.** Demographics.

| PID | Age | Gender | ECoG side | Moving Hand | Handedness |
| --- | --- | --- | --- | --- | --- |
| P1 | 76 | M | R | L | Both |
| P2 | 82 | F | R | L | R |
| P3 | 48 | M | R | L | R |
| P4 | 45 | F | L | R | R |
| P5 | 75 | M | L | R | Both |
| P6 | 73 | F | R | L | R |
| P7 | 71 | M | R | L | R |
| P8 | 63 | M | R | L | R |

**S1 Figure.**
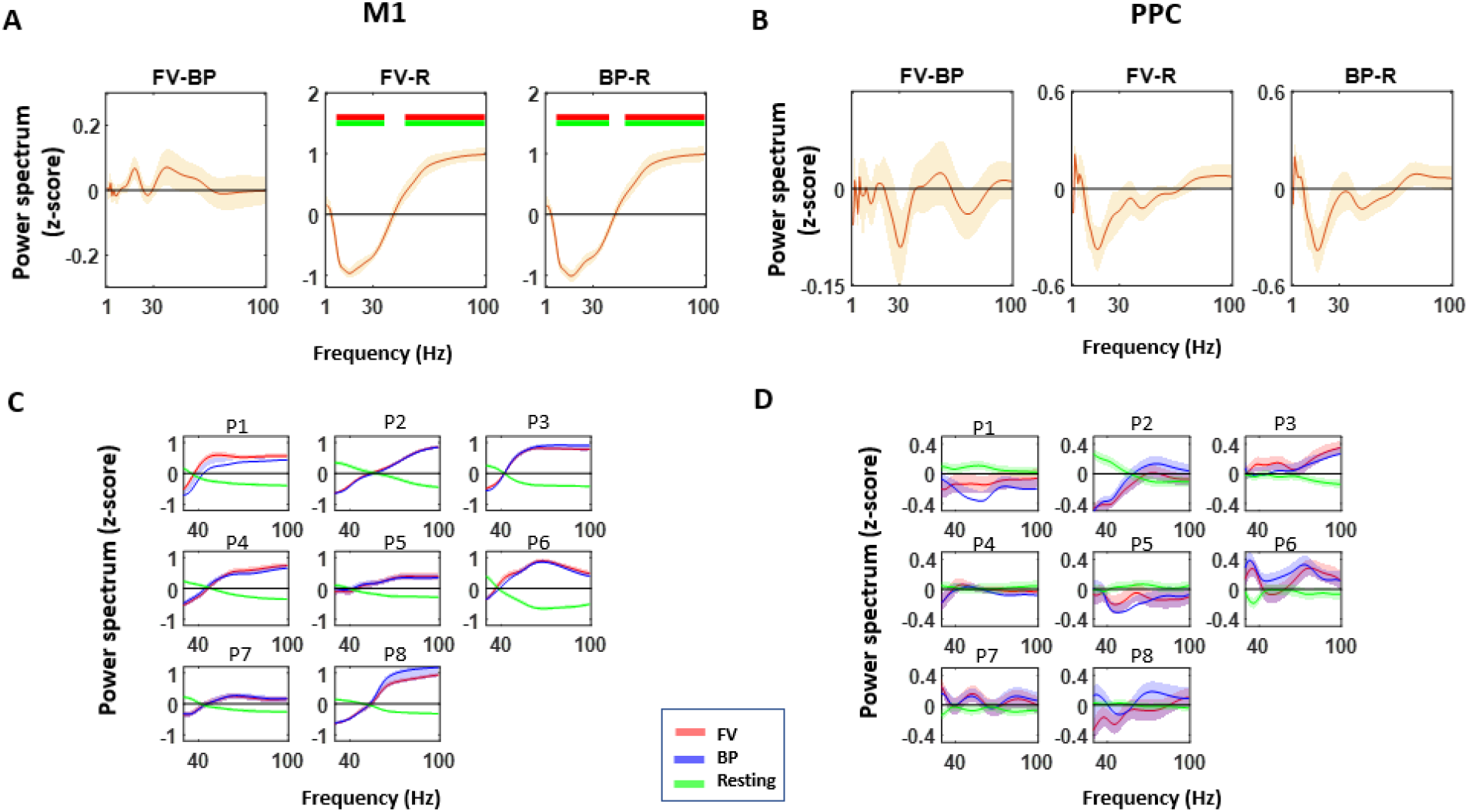
Oscillatory power does not differentiate between FV and BP movements in M1 and PPC. Power spectrum was computed per 1-second, and normalized (z-score) across time for each frequency (1-100Hz) using the BOSC toolbox (32), and plotted after averaging the z-scores per frequency and per movement type (red-FV, forward visually-guided; blue-BP, backward proprioceptively guided; green-rest). **A.** Normalized power spectral difference between FV and BP movements (left), FV and rest (middle), BP and rest (right) in M1, and **B.** in PPC. Data show mean values ± s.e.m. from nch = 8. Horizontal bars indicate significant differences using the Wilcoxen signed rank test for zero median (green-p<0.05, uncorrected; red-p<0.05/8 Bonferroni corrected) **C.** Normalized power spectrum for each participant (P1-P8) in M1, and **D.** in PPC. Shaded area represents s.e.m. of the normalized z-score per frequency. Gamma band is generally increased during movement within M1, but do not distinguish between FV and BP movements, and gamma band power does not characterize movement in PPC, nor distinguish the FV and BP movements.

**S2 Figure.**
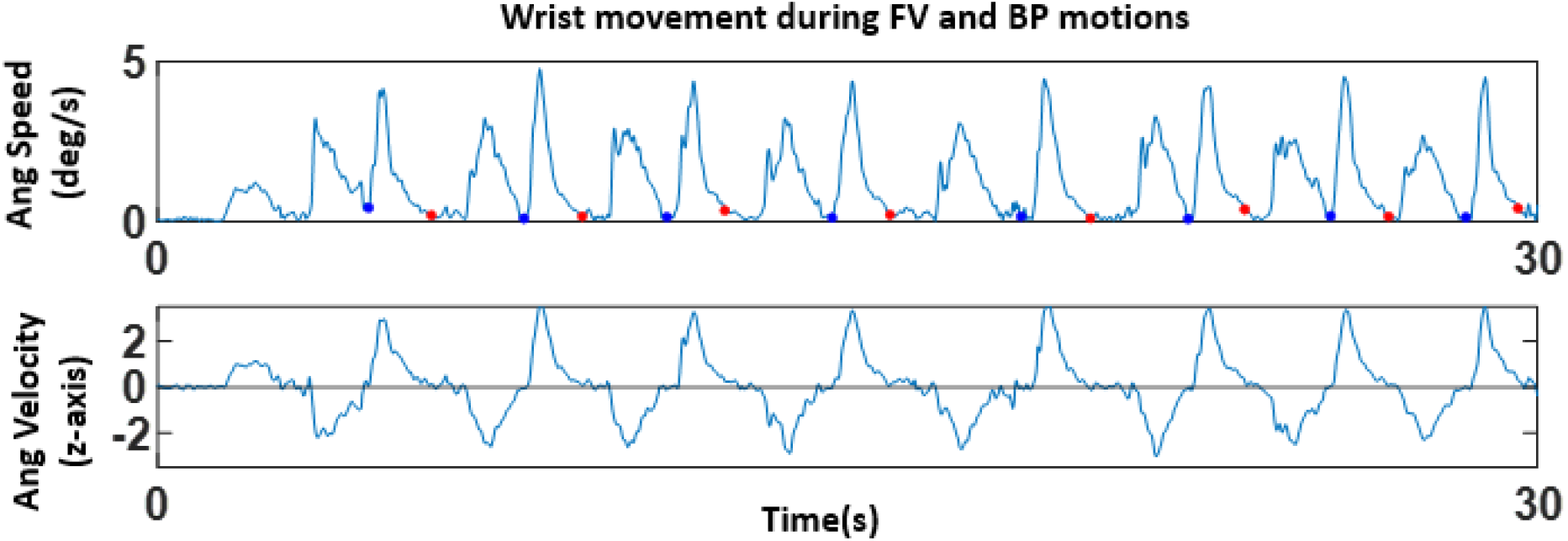
Kinematics during a representative participant’s reaching task. In order to distinguish FV and BP movements, angular speed (top) and angular velocity across z-axis (bottom) were examined together. The minima within angular speed (marked in blue and red) informed the time when the participant’s finger reached either the clinician’s finger or one’s own chin. The angular velocity zero-crossings informed of whether the goal was the finger or chin, such that when the value changed from negative to positive, this would indicate the chin; when the value changed from positive to negative, this indicated the finger reached the clinician’s finger.

**S3 Figure.**
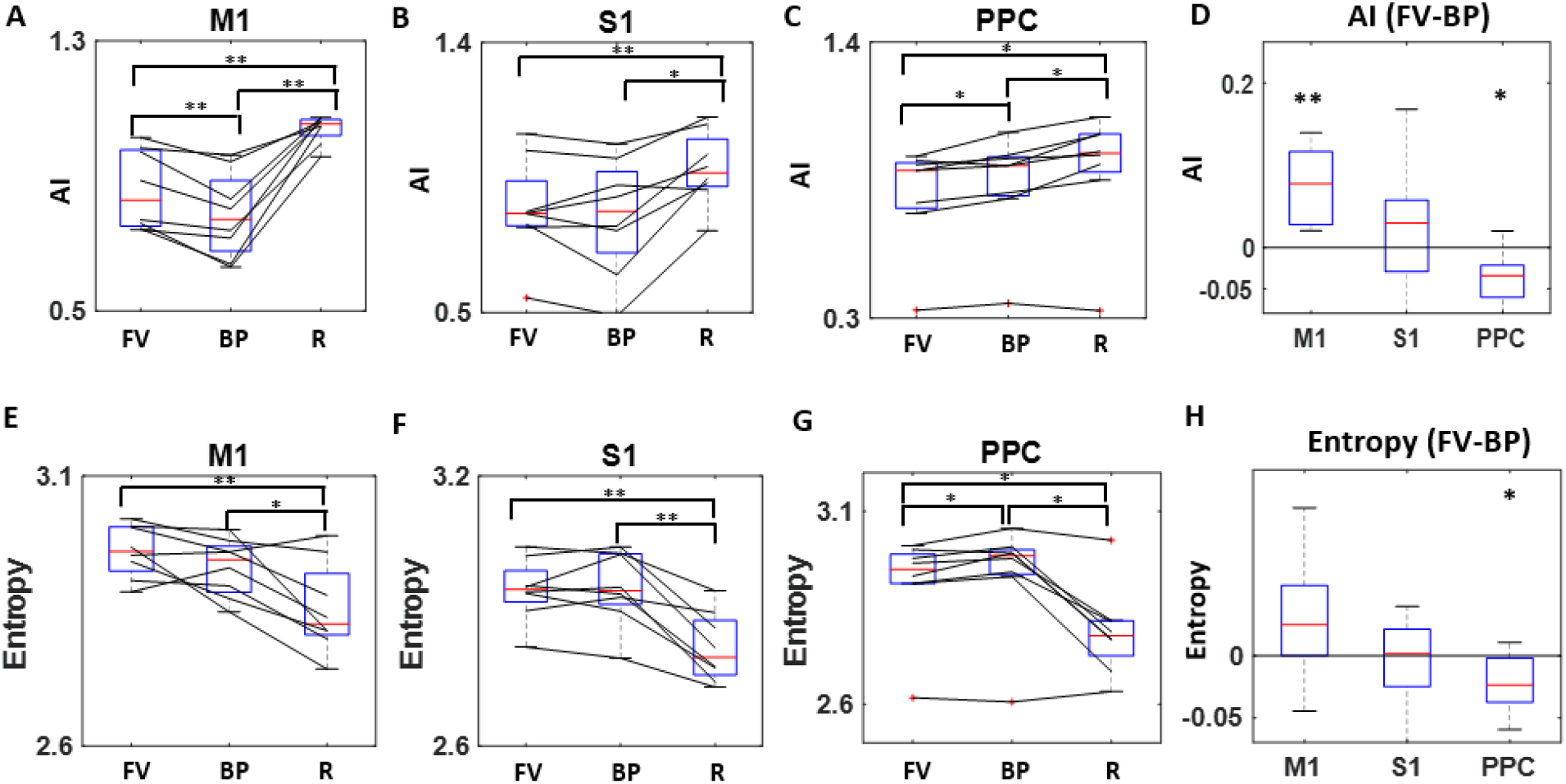
**A.** For exploratory purpose, we examined the AI when the 2 time windows were shifted by 20 ms, instead of 5ms shown in **Error! Reference source not found.**. We see a similar pattern of results with statistical pattern for M1, **B.** S1, and **C.** PPC, and **D.** their differentiation between FV and BP movements. **E.** Because entropy is an important feature of AI and TE metrics, we also examined the general entropy measure in M1, **F.** S1, and **G.** PPC, and **H.** their differentiation between FV and BP movements. Generally, we find the resting state (R) to show lowest entropy (i.e., most predictive) and this is in line with what we show in the Results. However, entropy is higher (i.e., more irregular/surprise) during FV movement in M1, and lower in PPC during BP movements, which is in contrast to what we show in the Results section. We interpret that the entropy measure we show here reflects an aggregate IEI stochasticity, whereas the analytics we introduce reflect the dynamical changes in the IEI’s. For that reason, we consider a general entropy metric to be capturing something different, but is beyond the scope of this study.

**S4 Figure.**
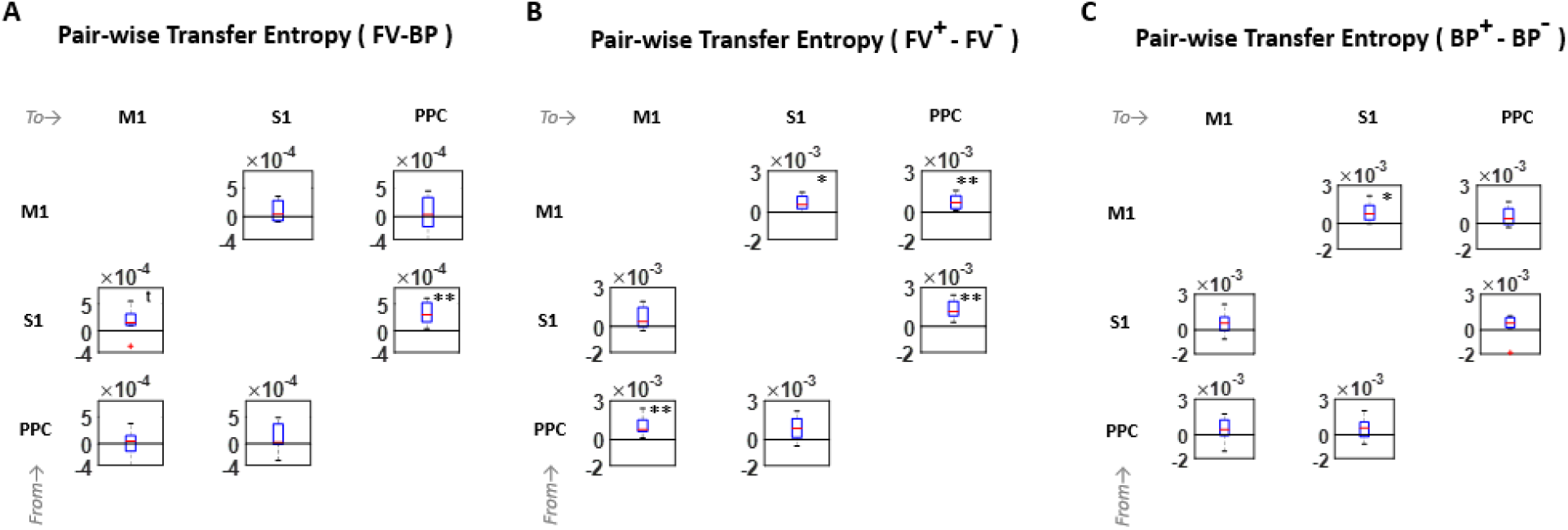
**A.** We examined the TE’s of all directional combinatorial pairs of the three cortical areas, and did a pairwise signed rank test to compare FV and BP movements. With the exception of S1 to M1, and S1 to PPC, all other directional paired flow do not distinguish the two movement types – FV and BP. **B.** When we further compare between accelerated FV (FV+) and decelerated FV (FV-) movements, generally there is a stronger connectivity during FV+ than FV-. Specifically, we found a stronger mutual connectivity between M1 and PPC, from S1 to PPC, and from M1 to S1. **C.** When we compared between accelerated BP (BP+) and decelerated BP (BP-) movements, the difference in connectivity was muted than during FV movement, with only significant difference from M1 to S1. *p<0.05, **p<0.01.

**S5 Figure.**
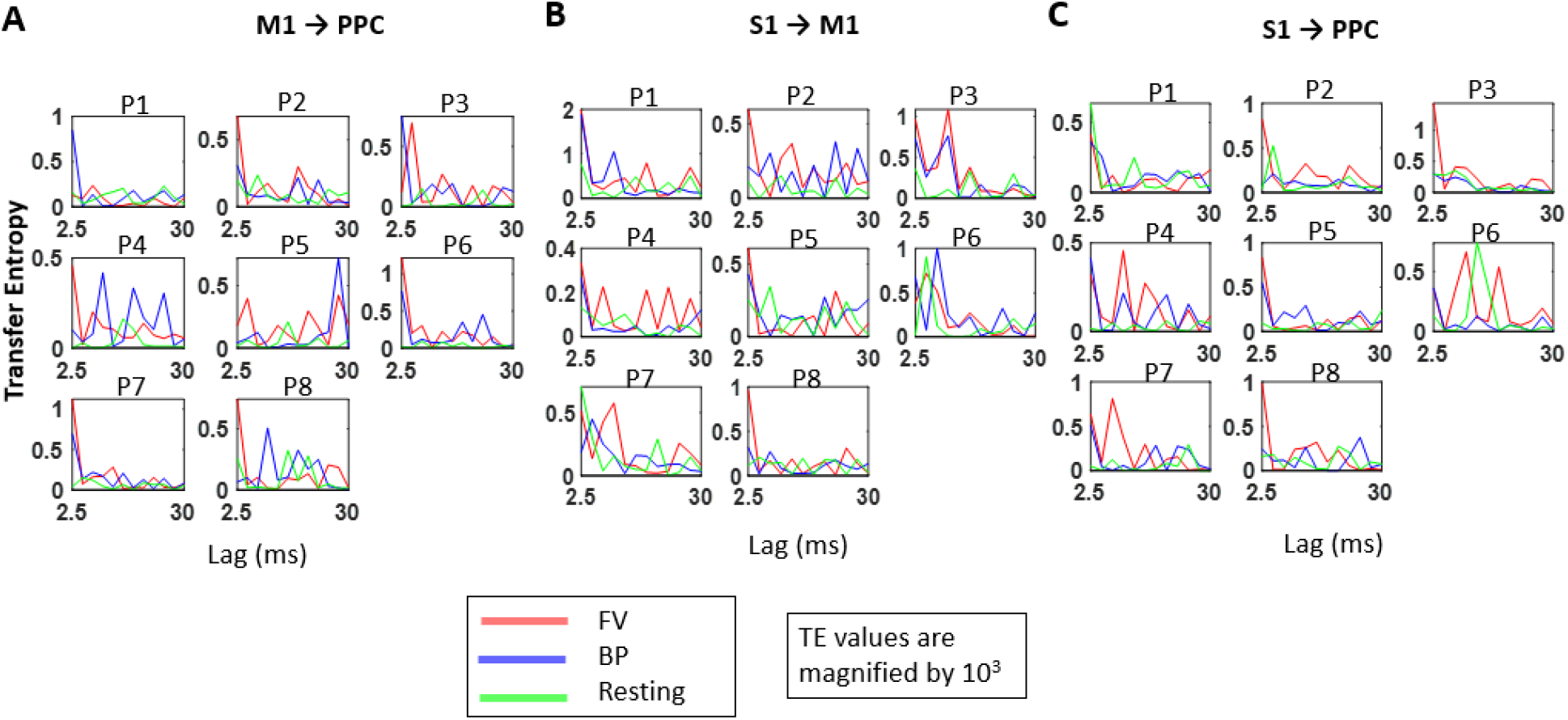
**A.** Transfer entropy at delays varying from 2.5 to 30ms for each participant (P1-P8) for M1 to PPC, **B.** S1 to M1, and **C.** S1 to PPC. In general the peak TE values occur early at around 2.5ms, but occasionally occur at around 10ms.

**S6 Figure.**
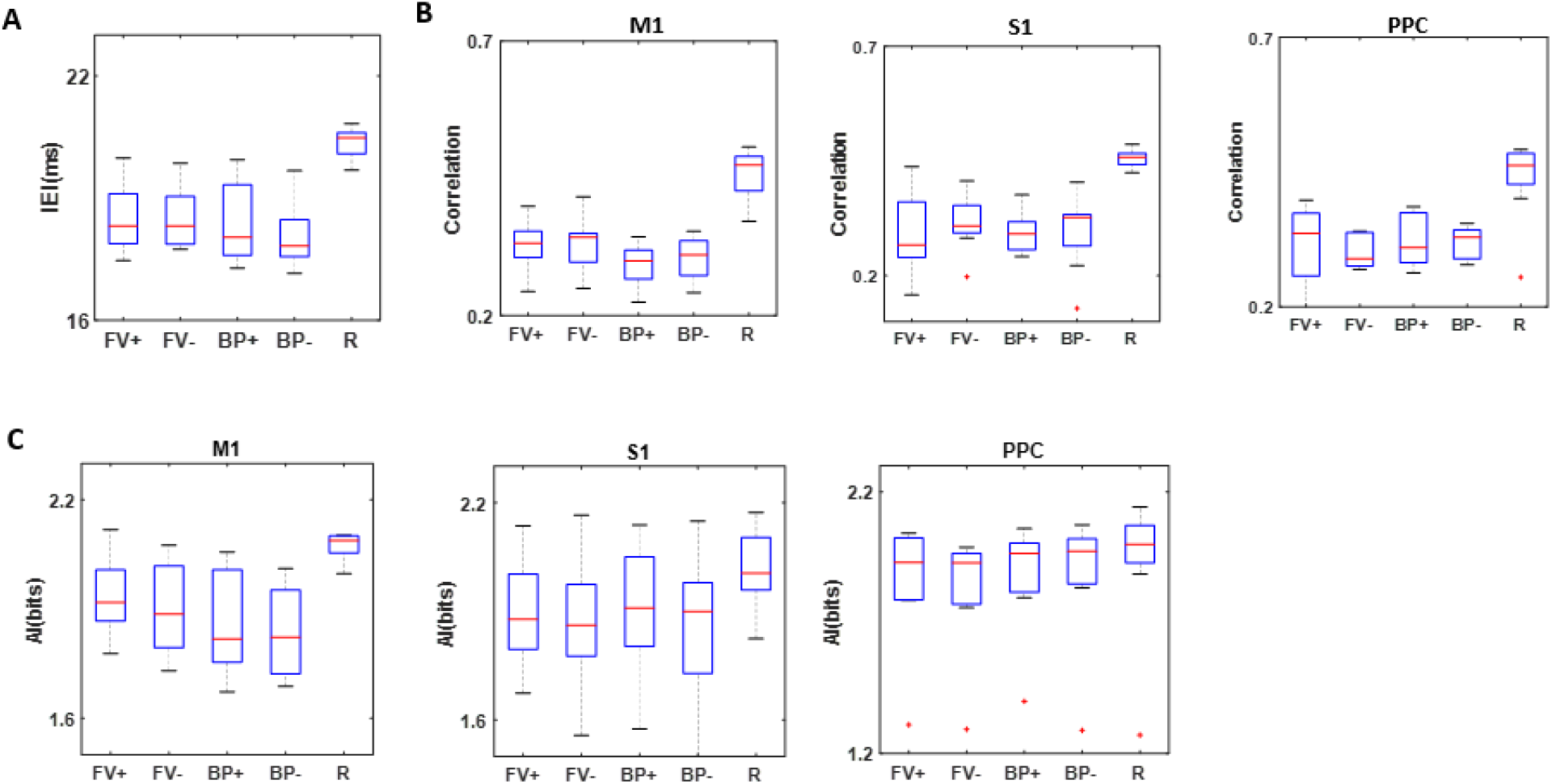
Irregularity of Gamma IEI does not differ between movements with different force dynamics. **A.** Mean IEI (ms) does not differ between accelerated FV (FV+) and decelerated FV (FV-), and does not differ between accelerated BP (BP+) and decelerated BP (BP-). **B.** Correlation between the Gamma cycle IEI and amplitude does not differ between FV+ and FV-, and between BP+ and BP-in M1 (left), S1 (middle), and in PPC (right). **C.** Auto-information (AI) does not differ between FV+ and FV-in M1 (left), S1 (middle), and PPC (right).

